# Formation of amyloid loops in brain tissues is controlled by the flexibility of protofibril chains

**DOI:** 10.1101/2022.09.13.507729

**Authors:** Alyssa M. Miller, Sarah Meehan, Christopher M. Dobson, Mark E. Welland, David Klenerman, Michele Vendruscolo, Francesco Simone Ruggeri, Tuomas P. J. Knowles

**Affiliations:** Yusuf Hamied Department of Chemistry, University of Cambridge, Cambridge, CB2 1EW, UK; Nanoscience Centre, University of Cambridge, Cambridge, CB3 0FF, UK; Laboratory of Organic Chemistry, Stippeneng 4, 6703 WE, Wageningen University & Research, the Netherlands; Physical Chemistry and Soft Matter, Stippeneng 4, 6703 WE, Wageningen University & Research, the Netherlands; Cavendish Labratory, Department of Physics, University of Cambridge, Cambridge, CB3 0HE, UK

## Abstract

Neurodegenerative diseases, such as Alzheimer’s Disease (AD), are associated with protein misfolding and aggregation into amyloid fibrils. Increasing evidence suggests that soluble, low molecular weight aggregates play a key role in disease-associated toxicity. Within these aggregates, protofibrillar loop-like structures have been observed for a variety of amyloid systems and their presence in brain tissues is associated with high levels of neuropathology. However, their mechanism of formation and relationship with mature fibrils has largely remained challenging to elucidate. Here, we use atomic force microscopy and statistical theory of biopolymers to characterise amyloid ring structures derived from the brains of AD patients. We analyse the bending fluctuations of protofibrils and show that the process of loop formation is governed by the mechanical properties of their chains. We conclude that *ex vivo* protofibril chains possess greater flexibility than that imparted by hydrogen-bonded networks characteristic of mature amyloid fibrils, such that they are able to form end-to-end connections. Furthermore, we show that these findings can be extended to several amyloid systems, giving a general framework relating the mechanical properties of assemblies and the conditions in which they can form loop structures. These results explain the diversity in the structures formed from protein aggregation and sheds light on the links between early forms of flexible ring-forming aggregates and their role in disease.

## Introduction

The study of protein aggregation and amyloid formation has garnered significant attention over the past few decades due to its role in human disease. This includes the aggregation of amyloid-β (Aβ) peptide, which is fundamentally implicated in the onset of Alzheimer’s disease (AD)^1,2^. Aggregation broadly involves the conversion of soluble monomers into insoluble amyloid fibrils which possess extended, highly ordered cross-β structure^1,2^. Current evidence points towards less organised, low molecular weight protofibrillar species as being the primary class of pathogenic structures^3–6^. Such species notably perturb healthy biological function, from disrupting cellular homeostasis by permeabilising biological membranes and non-specific receptor binding, to inducing inflammation and cell death^3,6,7^. This is consistent with the observation that soluble aggregates are associated with cognitive impairment in both mice and human models^8–10^.

Within this class of low molecular weight aggregates, ring-like loop structures have been reported for more than 10 different polypeptide systems^11^. This observation has resulted in much attention focusing on their disease-relevance and potential mechanisms of action. However, both the pathways by which they form as well as their relationship with mature fibrils has remained elusive. Much of our understanding comes from *in vitro* characterisation studies performed on recombinantly or synthetically-produced proteins^11–13^. Thus, it is unclear how these observations correlate with the properties of aggregates present in disease tissues. Recently, loop structures were reported in soluble fractions from the hippocampus of AD patient brains and characterised using a range of biochemical, biophysical and cellular assays^5^. These structures were associated with toxic effects, from increased levels of inflammation, membrane permeability, and cell death, suggesting that they do in fact play a role in disease pathology^7^. However, unravelling the specific conformation(*s*) related to disease is a key challenge.

Here, we employ single-molecule imaging and statistical theory of polymers to characterise the structure and mechanical properties of early AD brain-derived and *in vitro* protofibril species in both closed loop and open conformations, with the aim to understand the forces governing loop formation, and how these protofibrillar species compare to mature amyloid fibrils. To reach this objective, we measure the morphological and mechanical properties of brain-derived (ex vivo) pre-fibrillar aggregates by atomic force microscopy (AFM) and transmission electron microscopy (TEM). Then, additionally using amyloid assemblies formed by bovine α–lactalbumin as a model system, we measure the arc–length distribution of filament and loop structures and show that a theory of semi–flexible polymers quantitatively predicts the range over which the formation of closed rings is expected to be favoured over open protofibril chains. Finally, we demonstrate that this type of model can be extended to other systems and provides a general framework which links the mechanical properties with the morphology and toxicity of amyloid structures.

## Results

### Characterisation of amyloid loops

We first induced the formation of amyloid protofibrils and rings from recombinant α–lactalbumin *in vitro* (Fig. 1 A) Conditions were selected to destabilise the native state of the protein, namely low pH (2.0) and high temperature (50°C). Initially non-fibrillar aggregates were observed, followed by the formation of chain–like aggregates after 2 h of incubation (Fig. S1 A). Amyloid loops and open chain protofibrils were identified after incubation times ranging from 5 to 14 h (Fig. S1 B and C). After prolonged incubation at high temperature (72 h, Fig. S1 D) the filaments and loops were no longer observed, and only aggregates of smaller size were seen, suggesting that the flexible chains possess a low level of structural robustness, an observation in agreement with the mechanical characterisation presented below. Decreasing the incubation temperature slowed the kinetics of amyloid fibril formation by bovine α–lactalbumin, consistent with the presence of a free energy barrier resulting from the structural reorganisation required associated with the conversion of the protein to the amyloid form. For example, amyloid rings were observed after 1 day of incubation at 37°C and after 1 month of incubation at −5°C. This contains information on the combined nucleation and growth rates, giving an order of magnitude estimate which is comparable to that measured for the growth of insulin fibrils^14^. For the sake of simplicity, protofibrils formed from α–lactalbumin will henceforth be referred to as ‘in *vitro*’ samples.

**Figure 1.**
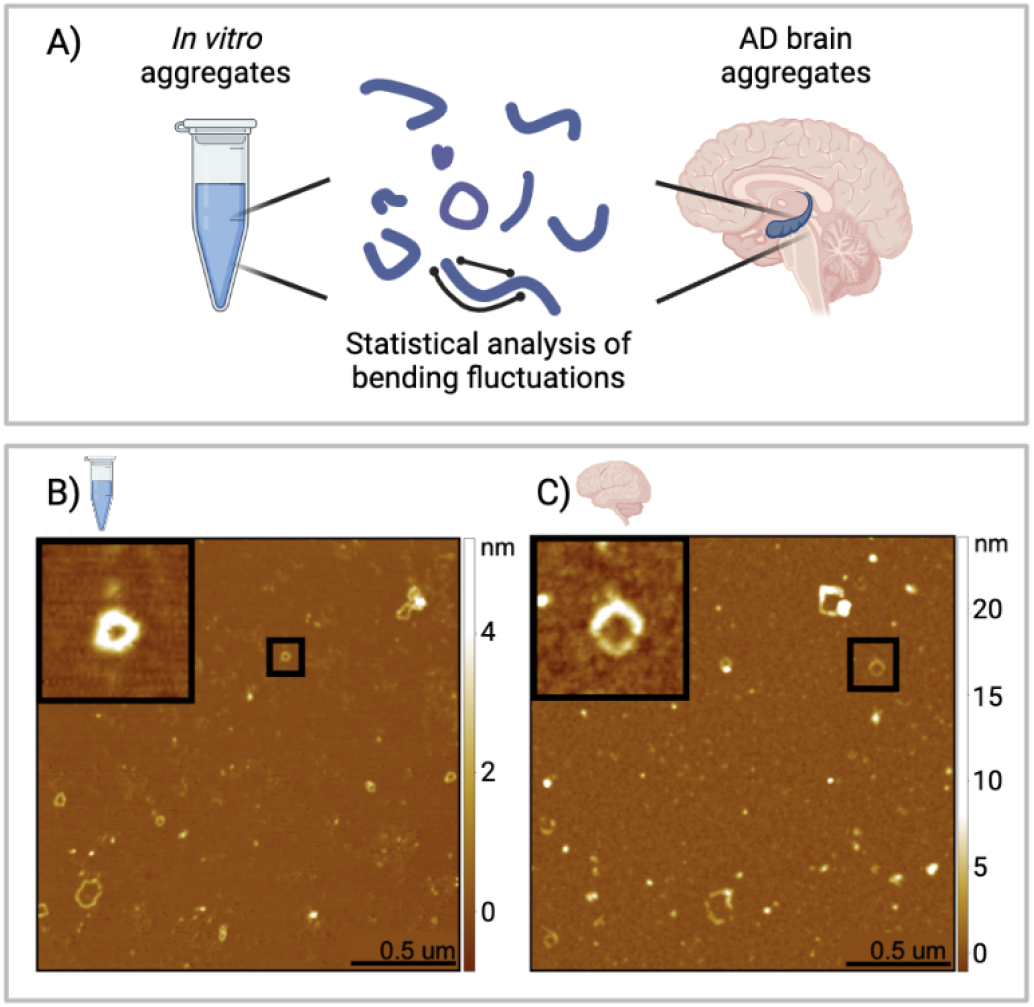
AFM images of *in vitro* and *ex vivo* protofibrillar loops. (A) Protofibrillar aggregates were extracted and studied by statistical analysis of their bending fluctuations. Samples analysed were recombinant α-lactalbumin protofibrillar aggregates (in *vitro*) and soluble aggregates extracted from the hippocampus of AD patient brains (ex vivo). (B) Example AFM images are shown for in *vitro* and (C) ex vivo samples. The black squares show a zoom of examples of closed ring structures.

We then extracted and characterised soluble aggregates from the hippocampus of post-mortem Alzheimer’s disease brains. Briefly, brain homogenate was subjected to successive rounds of buffer soaking and centrifugation, then the soluble fraction was retrieved and deposited on a mica surface and characterised using AFM^5,15^. We observe heterogeneous structures, including spherical oligomers and protofibrillar structures which can exist in both a closed loop and an open conformation, similar to those observed in the *in vitro* system (Fig. 1B).

### Mechanical properties of amyloid loops

Having established the presence of ring-like structures for the *in vitro* and *ex vivo* samples, we sought to define their length distribution. Over 1000 individual structures from over 200 AFM maps were individually traced (Fig. 2) using a semi–automatic algorithm (Materials and methods), and the arc–lengths were measured from the digitized contours using fitted splines. Additionally, the height was measured for the same structures by considering the maximal height from the local average background from sections perpendicular to the tangent of the spline following the contour of the filament (Fig. S2). These measurements allow us to understand the mechanical properties of protofibrils, such as the persistence length *l_p_* which can be calculated directly from AFM maps using the mean square end-to-end distance *R* as a function of contour length *L* (Fig. 2E):

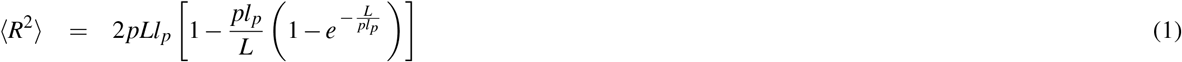

*p* is a surface parameter, which has a value of 2 for chains that have equilibrated on the 2D surface and 1.5 ± 0.5 for those that have not. This value can be determined based on the scaling exponent of the plot of R versus contour length, and was found to be 0.74 and 0.79 for endogenous and synthetic protofibrils, respectively, indicating they have equilibrated in 2D^16,17^.

**Figure 2.**
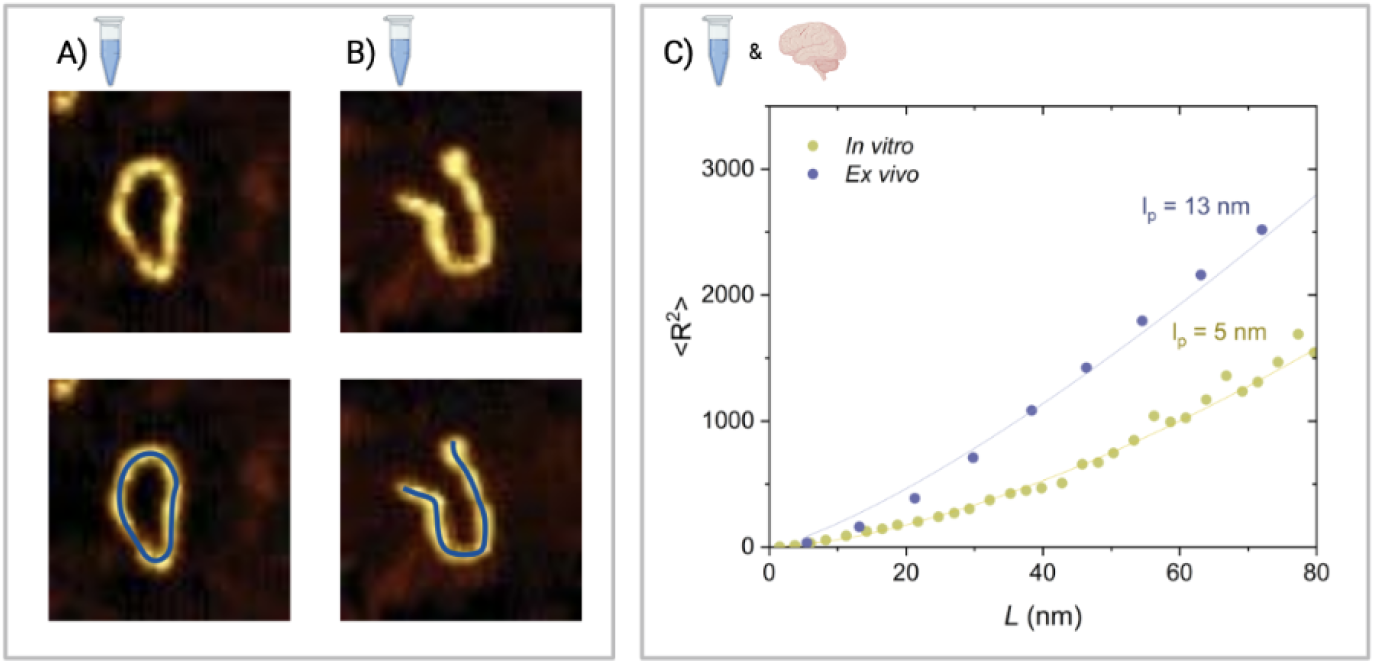
Measuring protofibrils to calculate the persistence length. (A) Open and (B) closed rings are observed for α–lactalbumin. (C & D) Protofibrils were traced using a semi-automated algorithm by fitting along the protofibril spline (blue line). (E) Persistence length *l_p_* for in *vitro* (n=426, 161 images) and ex vivo (n=24, 41 images) samples can be calculated based on the plot of 〈R^2^〉 versus the contour length L, as measured from the AFM maps.

From equation 1, we estimated the *l_p_* values for in *vitro* and ex vivo protofibrils. In the case of in *vitro* protofibrils, the uniform nature of the sample allows us to determine a single value of *l_p_* of ~·5nm. For the ex vivo aggregates, the heterogeneity of human samples means that one value of lp does not accurately represent the sample. As such, we performed a grouped analysis of the protofibrillar aggregates as a function of their height, which is indicative of their aggregation maturity^18,19^. The groups we selected are 0-3nm, 3-6nm, 6-9nm, and 9-12nm, which corresponds with the height of protofilament chains (Fig. S3). The majority of ring-like structures were found in the 3-6 nm height group, with an *l_p_* of ~13 nm. We used this group as representative of the flexible ex vivo samples as it corresponds well with the height observed for in vitro protofilaments, facilitating comparison. We next calculated the bending rigidity (CB) from the values of *l_p_*, which were found to be 5.1 x 10^-28^ N/m for ex vivo protofibrils and 2.0 x 10^-28^ N/m for in *vitro* protofibrils. These values are significantly lower than those reported for mature amyloid fibrils,^20^, indicating a lack of intermolecular hydrogen-bonding and extended cross–β structure^19,21,22^.

### Modelling the probability of loop closure

Having characterised the mechanical properties of protofibrils, we next sought to understand the likelihood of finding a protofibril in a closed or open conformation. We measured the incidence of closure as a function of the arc–length. The data in Fig. 3 shows that for small values of the arc–length, all protofibrils are found as open chains; with increasing arc–length, the probability of finding loops increases, and finally for very long chains decreases again. In order to interpret these measurements in a quantitative way, we consider a simple model based on statistical theory of biopolymers. At any given time a fibril composed of *j* monomers can exist either as an open chain configuration with two free ends, or in a closed loop state where the ends are joined. These two forms are inter–converted with rates that in general depend on the length of the chains: 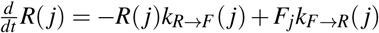 where *F_j_* and *R_j_* are the concentrations of the open and closed forms, and *k_F→R_*(*j*) and *k_R→F_* (*j*) are the first order rates for ring closure and opening.

**Figure 3.**
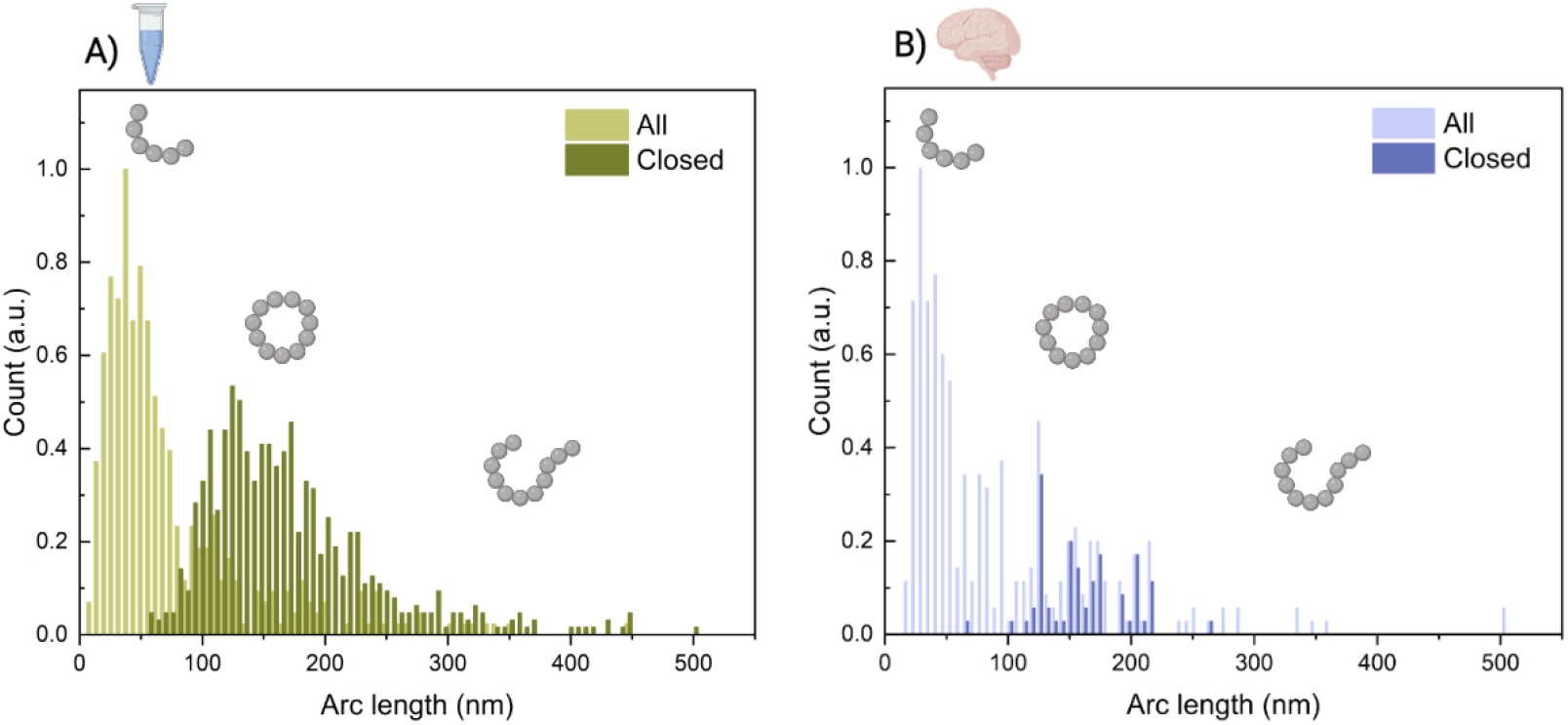
Arc length distributions of both open and closed conformations. Distributions of both open and closed conformations were measured for (A) in *vitro* (n=1022, 161 images) and (B) *ex vivo* (n=324,41 images) protofibrils.

We first consider the ring closure rate: 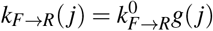 whereby the *j* dependence is in the circularisation probability *g*(*j*), and the constant factor 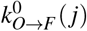 contains information about the diffusion coefficient of the ends of the polymer as well as about the reaction cross-sections. The circularisation probability for semi-flexible chains has been explored theoretically by Stockmayer and Yamakawa^23^ and more recently by Liverpool and Edwards^24^ and is very generally given by the sum of the Boltzmann weights corresponding to different configurations of the bond vectors 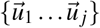 which satisfy the loop closure condition 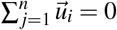:

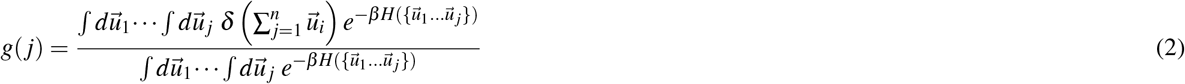

with the semi-flexible Hamiltonian: 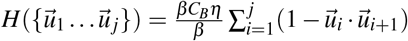 We take here the result based on perturbation theory derived by^23^, which enables an approximate value of the circularisation probability to be derived for semi-flexible chains in the continuum limit; the result can be expressed as a simple function of a scaled dimensionless arc length, 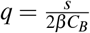:

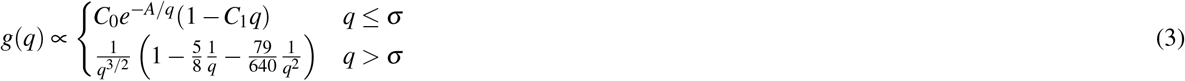

where the approximate numeric values of the dimensionless constants are σ = 0.96, C_0_ = 1.51 · 10^3^, C_1_ = −0.81 and *A =* 7.026.

We furthermore include the possibility for the two ends of the chain to break apart again, and this process can be modelled as a thermally activated barrier crossing, where in the general case the barrier height depends on the strain induced by the ring closure condition: 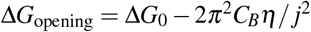 where Δ*G*_0_ is the free energy barrier in the absence of strain, *C_B_* is the bending rigidity of the chain, and *η* is the number of monomers per unit length. Therefore the opening rate can be written as: 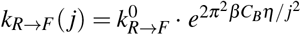 where 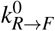 is a constant which contains information on the absolute value of Δ*G* and 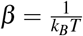 is the inverse temperature.

If the system is in a steady state with respect to the conversion between open ended filaments and loops, the probability *P*(*s*) of finding a given structure in the closed configuration as a function of the arc–length s is given by: *P*(*j*) = *r*(*s*) /(*r*(*s*) + 1) where *r*(*s*) is the ratio of the opening and closing rates (Fig. 4):

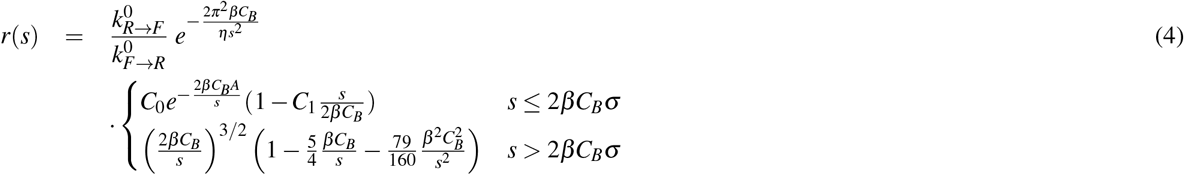

**Figure 4.**
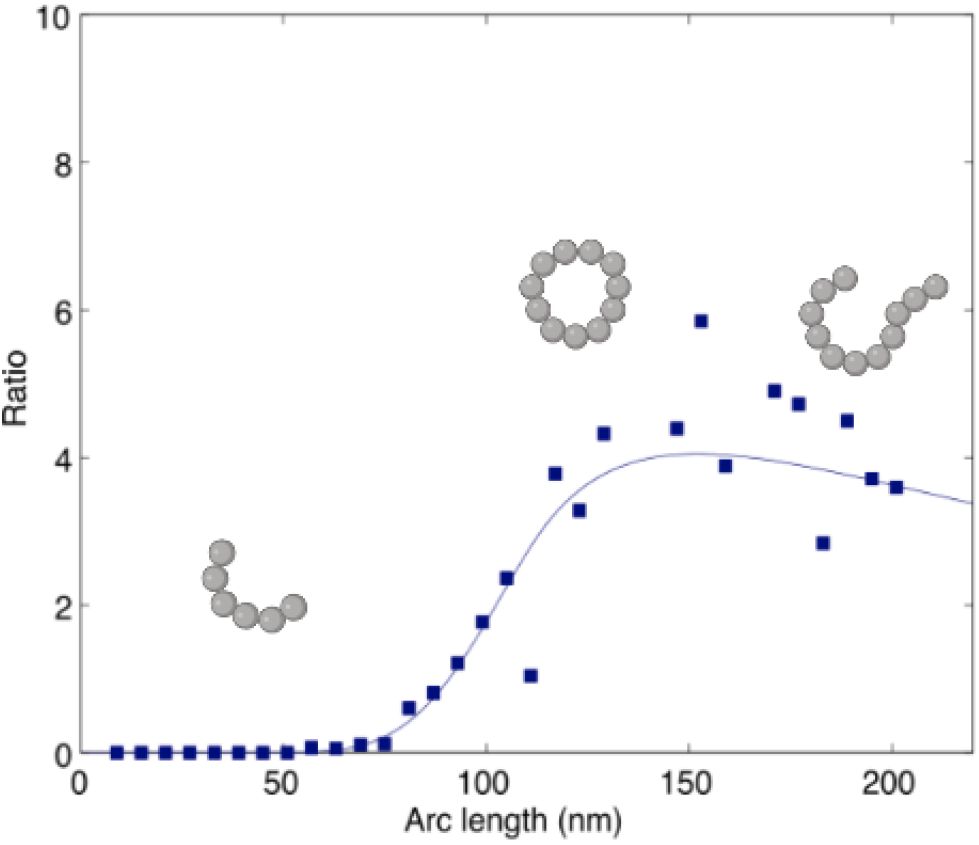
Modelling the probability of loop closure as a function of the arc length. The probability of loop closure was modelled as a function of protofibril length, based on theory of semi-flexible polymers. The y-axis reports the ratio of the opening and closing rates. From equation 4 (main text), as the arc length increases, so does the likelihood of finding a protofibril in a closed loop conformation. Above a certain length, the likelihood decreases again as the probability of two open chain ends finding eachother decreases.

A two parameter fit to the measurements in Fig. 3 of Eq. 4 where the unit–less pre–factor 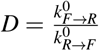 and the bending rigidity *C_B_* are left free yields the value *D* = 41.7 and *C_B_* = 2.4 · 10^-28^ N m^2^, which is consistent with the value obtained by statistical analysis of AFM images. This value of the bending rigidity controlling the loop formation is also of a similar order of magnitude as that found for the open chains of α-lactalbumin of *C_B_* = 1.4 · 10^-28^ Nm^2^ from an independent analysis of shape fluctuations^14^, showing that the intrinsic mechanical properties of the filaments determine the distribution of ring structures. The values of the bending rigidity reported here are also below that which is typically measured for elongated amyloid fibrils, suggesting that the *in vitro* and *ex vivo* protofibrils have a lower level of structural organisation than that which is generally found in amyloid fibrils.

The present measurements also enable us to investigate the energetics of the ring closure. There is an enthalpic energy cost of Δ*H* = 2*C_B_π*/*d* > 0 to bring the two ends of the chain together, and therefore the energy gain from the ligation of the ends must exceed this value for this state to be significantly populated. We observe loops down to a diameter of approximately *d* = 19 nm, and therefore the interaction between the filament ends must contribute at least 11 kcal/mol, an energy which is equivalent to that gained from the formation of 2–3 hydrogen bonds. The mechanical strain within the smallest loops can be estimated from geometric considerations: *ε* = Δ*l*/*l*_0_ = *h*/*d* ≈ 5.2% where *h* is the height of the filaments, and *d* the diameter of the ring. This value is smaller than the maximal strain which the cross-β core structure in insulin fibrils was shown to be able to accommodate without fracturing (18 %)^25^.

Finally we can test the generality of the model by considering the measurements of the rigidities of other amyloid fibrils. A plot of the persistence length vs. the arc length (Fig. 5) for different fibril systems reveals that the structures which have a ratio of persistence length to arc length less or equal to one exist in the ring configuration, whereas structures with a ratio significantly above one only exist as linear fibers.

**Figure 5.**
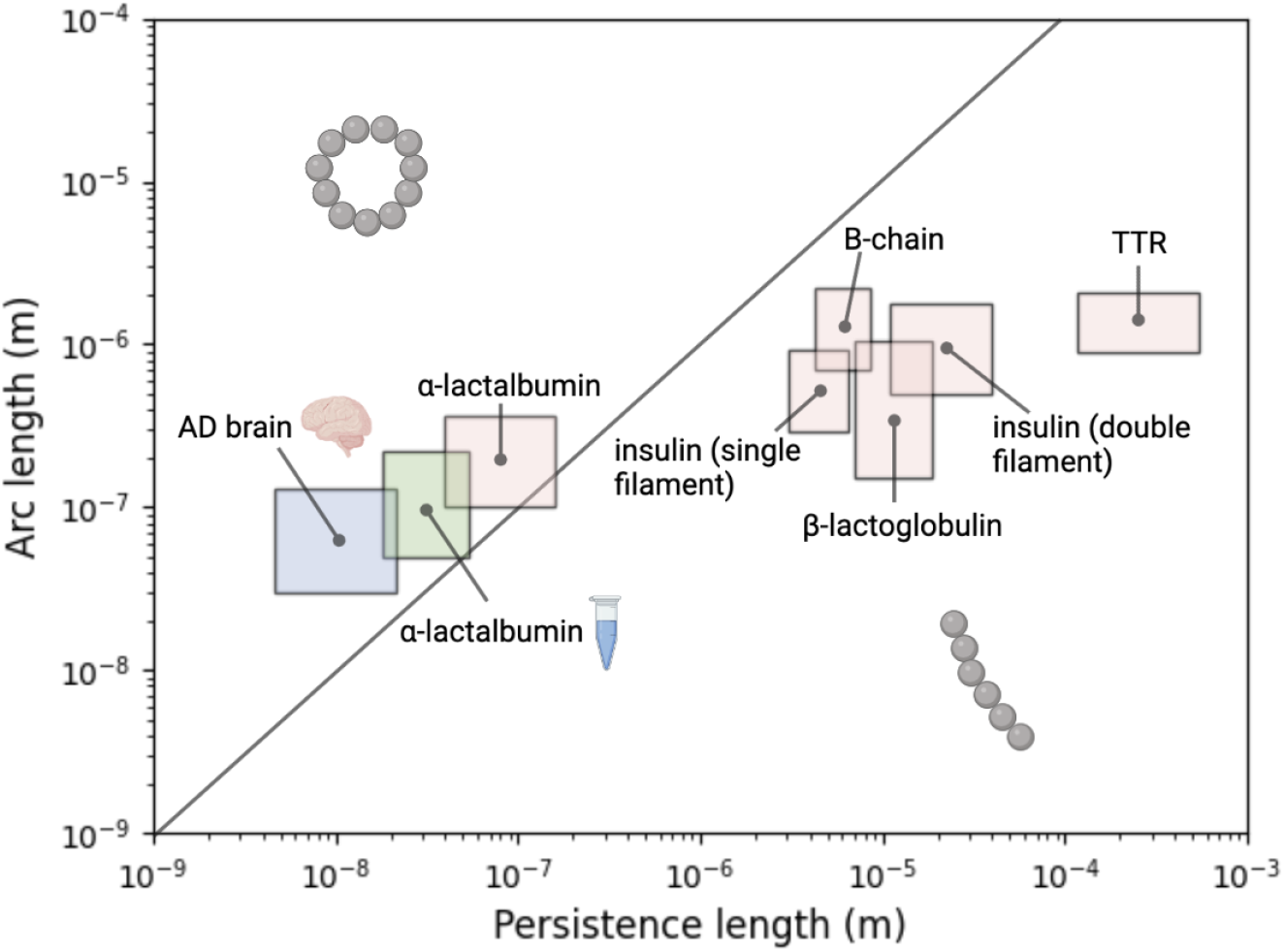
Persistence length plotted as a function of arc-length for a variety of amyloid protofibril and fibril assemblies. The persistence length and arc length are plotted for a variety of amyloid systems. From the present work are ex vivo aggregates and in *vitro* (α-lactalbumin) aggregates, incdicated by the brain and test tube, respectively. Literature values are plotted for 1) α-lactalbumin, 2) insulin (single filament), 3) B-chain, 4) *β*-lactoglobulin, 5) insulin (double filament), and 6) TTR from ref^14^. Where the ratio between persistence length and arc length is below one (indicated by black line), loop structures may form.

## Discussion

We have described the mechanical properties of brain-derived protofibrils and the forces that govern their loop formation in order to increase our understanding of the nature of loop structures in neurodegenerative disease. We have demonstrated that protofibrillar structures do not possess the extended hydrogen-bonded internal structure of mature amyloids, such that they are capable of forming and breaking end-to-end connections. The likelihood of finding a protofibril in a closed loop can be simply understood as the probability of the two ends finding eachother. Previously reported *in vitro* loop structures were more homogeneous in size, leading researchers on the search for a uniform, rigid class of structures, such as a pore-forming hexame^r12,13,26^. This is inconsistent with the data presented here on brain-derived aggregates, which are heterogeneous both in morphology and whether they are found in a loop or stretched-out conformation. This adds evidence to the idea that the search for a single toxic species in AD may be a fruitless one, and that rather a heterogeneous, dynamic population of low molecular weight structures are involved in disease pathogenesis^6^.

While we have not investigated the mechanism of action of soluble protofibrils, this has been investigated in previous work, making it possible to draw links between the mechanical properties established here and their biological function^5^. It is clear that these structures capable of forming loops are distinct from mature amyloid fibrils found in plaques of AD brains, not only in size and solubility, but also in internal structural properties. These differences in mechanical properties not only explain the ability certain amyloid species possess to form loops, but may also explain the discrepancies in mechanisms of actions between mature and earlier, protofibrillar structures. These soluble, low molecular weight species were associated with increased levels of inflammation, membrane permeability and cell death, suggesting that these are key disease-relevant structures^5^. This toxicity may be imparted by the low level of structural robustness, such that they are more capable of forming non-specific, aberrant interactions with cell membranes and receptors.

## Materials & Methods

### Preparation of endogenous aggregates from human brains

Endogenous aggregates were extracted from Alzheimer’s disease as previously described^5^. Briefly, these experiments employ aggregates derived from fresh frozen brains from three Alzheimer’s disease patients diagnosed as being at Braak stage III. Soluble aggregates were obtained by following a previously established protocol with a few adaptations^4^. Human brain tissue was chopped into 300 mg pieces using a razor blade and incubated with gentle agitation in 1.5 mL of artificial cerebrospinal fluid (aCSF) buffer (124 mM NaCl, 2.8 mM KCl, 1.25 mM *NaH_2_PO_4_*, 26 mM *NaHCO_3_*); pH 7.4, supplemented with 5 mM EDTA, 1 mM EGTA, 5 μg/mL leupeptin, 5 μg/mL aprotinin, 2 g/mL pepstatin, 20 μg/mL Pefabloc, 5 mM NaF) at 4°C for 30 min. Samples were centrifuged at 2,000 g at 4°C for 10 min and the upper 90% of the supernatant was collected and centrifuged at 14,000 g for 110 min at 4°C. The upper 90% of the supernatant was extracted and dialysed using Slide-A-Lyzer™ cassettes (Thermo Scientific, Cat. 66330) with a 2 kDa molecular weight cut off, against 100-fold excess of fresh aCSF buffer with gentle agitation at 4°C. Buffer was changed three times over the course of 72 hours dialysis. The prep was carried out under sterile conditions, using autoclaved LoBind Eppendorf tubes and pre-sterilised pipette tips to reduce endotoxin contamination. Samples were aliquoted into small volumes, snap frozen and stored in a −80 ^°^C freezer and thawed only once prior to experimentation.

### Formation of amyloid fibrils by bovine α-lactalbumin

α–Lactalbumin from bovine milk (Type III, calcium depleted, Sigma-Aldrich) was dissolved at 5 mg/ml in *H_2_O* adjusted to pH 2.0 with HCl, with 20 mM dithiolthreitol (DTT), and incubated at 50°C. After 3-6 hours incubation at 50°C, amyloid fibrils, both in the form of open chains and loops were identified by TEM, and further confirmed by a positive interaction with the dyes Thioflavin-T and Congo Red. Amyloid fibril formation preceded more slowly at lower incubation temperatures: amyloid loops were visible by TEM after 1 days incubation at 37°C, after one month at 5°C.

### Transmission electron microscopy (TEM)

Transmission electron microscopy (TEM) Nickel electron microscopy grids with carbon–coated formvar support films were prepared for microscopy by the addition of 21 of protein sample at a concentration of 1 mg/ml. The grids were then washed with 3 x 10 μl *H_2_O* and negatively stained with 10 μl of uranyl acetate (2 % (w/v), Agar Scientific). Samples were viewed under 20125-K magnifications at 120 kV acceleration voltages using a Philips CM100 transmission electron microscope.

### Atomic force microscopy (AFM)

α-lactalbumin protein solution (0.05 – 1.0 mg/ml) was deposited onto freshly cleaved mica substrates (Agar Scientific Ltd., Stansted, Essex) and air-dried. Topographic data were then acquired with a Dimension 3100 SPM (Veeco Instruments Inc., Woodbury, NY, USA) in conjunction with a Nanoscope IV control system operating in tapping mode. Ultrasharp Micromasch silicon cantilevers (NSC12/Si3N4/50) were used at resonance frequencies between 150 kHz and 225 kHz.

Endogenous protein imaging was performed in a similar way. Protein solutions were diluted 10x in PBS buffer, and 10 μl was deposited onto freshly cleaved mica. The samples were incubated for 10 min, followed by rinsing with 1 mL milliQ water. The samples were then dried using a gentle flow of nitrogen gas. AFM maps of 3-D morphology were acquired in regime of constant phase change, with 2-4 nm/pixel resolution using a NX10 (Park Systems, South Korea) operating in non-contact mode. This set up was equipped with a silicon tip with a nominal radius of <10 nm and spring constant of 5 N/m (PPP-NCHR). Scanning Probe Image Processor (SPIP) (version 6.7.3, Image Metrology, Denmark) software was used for image flattening and single aggregate statistical analysis. The average level of noise for each image was measured using SPIP software and was smaller than 0.1 nm. All the measurements were performed at room temperature.

### Statistical Analysis

The arc length and shape of protofibrils were measured using a semi-automated algorithm that extracts the highest points of the protofibril axis and fits a 2-dimensional parametric spline to these points. The *lp* values were then extracted from statistical analysis of these shape fluctuations, as described in the main text, using DNA Trace software^27^. Height of protofibrils were measured by considering the average maximal height from the average height of the background, and was found to be 2.2 ± 1.3 nm for endogenous aggregates and 1.9 ± 0.4 for synthetic aggregates.

## Acknowledgements

We wish to acknowledge funding support from the ERC, EPSRC and Newman Foundation. We also thank the Cambridge Brain Bank and Addenbrooke’s hospital, Cambridge, for processing and providing Alzheimer’s disease brain tissue for this study. The Human Research Tissue Bank is supported by the NIHR Cambridge Biomedical Research Centre.

