## Supplementary Info for "Formation of amyloid loops in brain tissues is controlled by the flexibility of protofibril chains"

July 13, 2022

### Supplementary figures

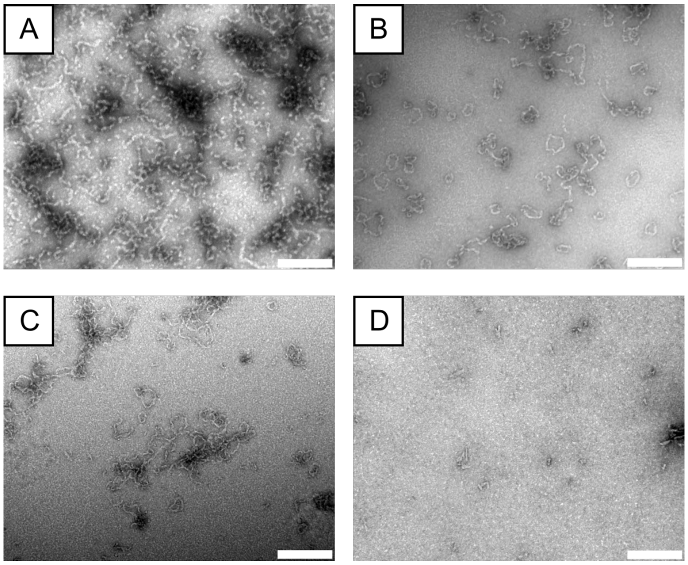


**Figure 1. TEM imaging of recombinant $\alpha$-lactalbumin aggregates**. (A-D) $\alpha$-lactalbumin was imaged via TEM in destabilising conditions after A) 2 (A), 5 (B), 14 (C), and 72 (D) hours incubation time. Protofibrillar structures are initially observed (A), followed by the observation of closed ring structures (B & C). After prolonged incubation (D), the elongated structures are no longer observed.

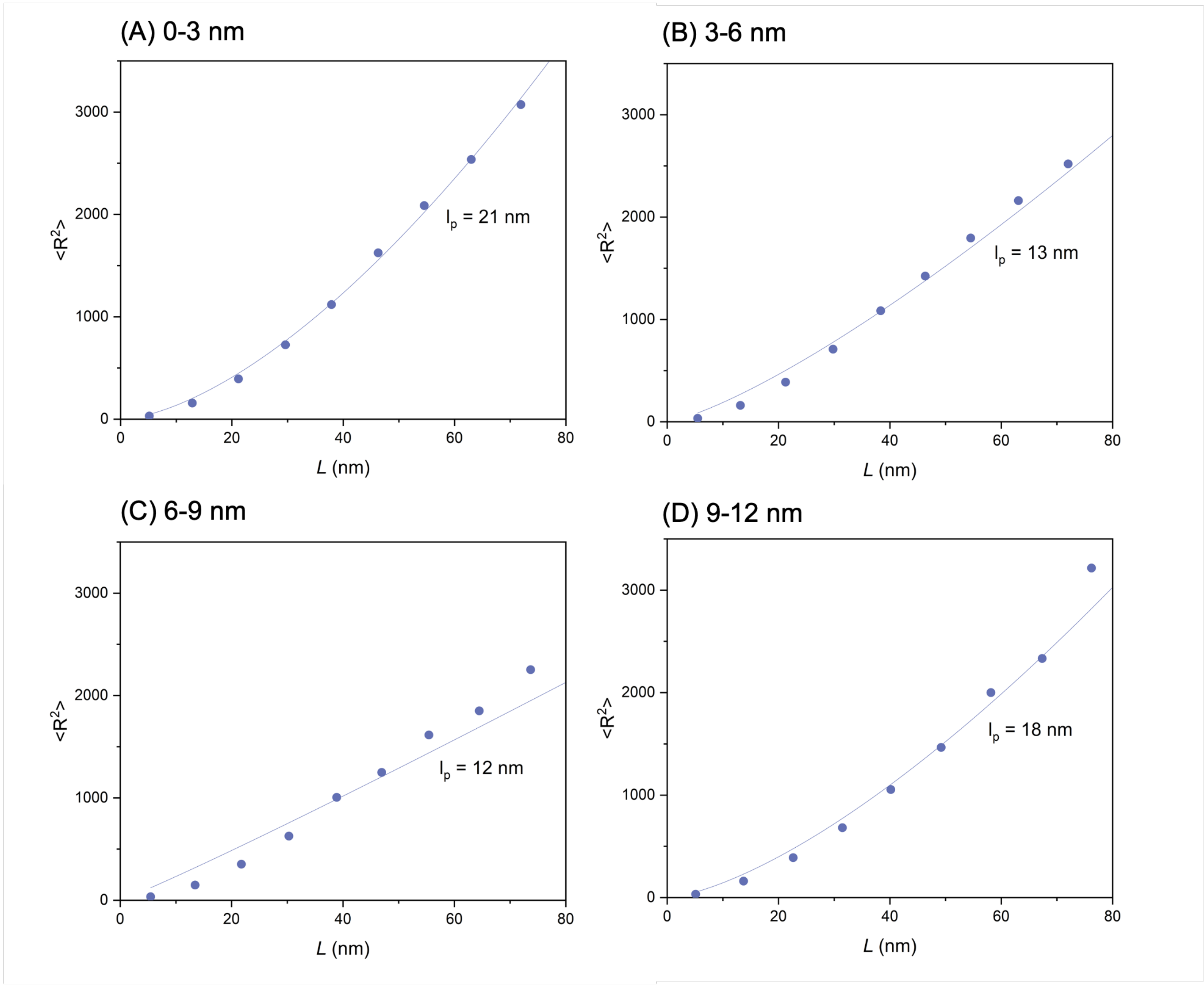


**Figure 2.** Plots of contour length ($L$) vs $R^2$ for *ex vivo* protofibrils. Due to the heterogeneous nature of brain samples, it is likely that more than one protein species exists. Analysis was therefore performed in groups corresponding the height of 1 (n=137), 2 (n=24), 3 (n=28), and 4 (n=9) individual protofilaments.

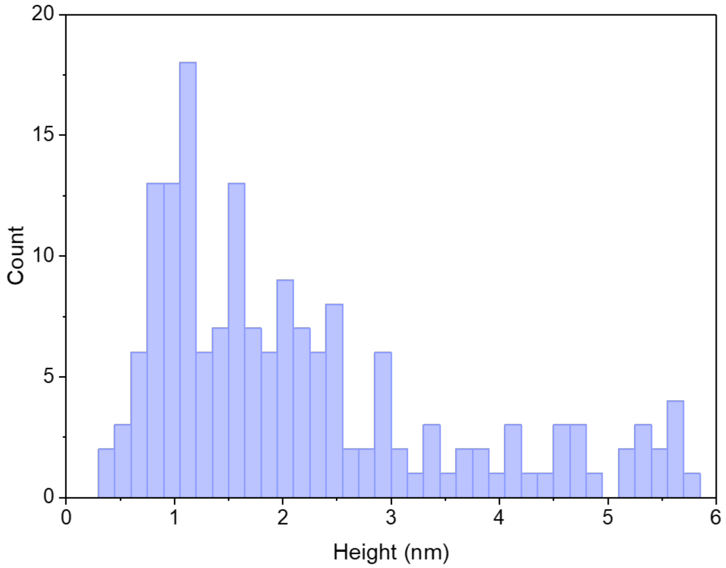


**Figure 3.** The cross-sectional height distribution for *ex vivo* aggregates was measured using AFM. The mean height was 2.2 ± 1.3 nm (n=198).
